# Macromolecule Translocation in a Nanopore: Center of Mass Drift–Diffusion over an Entropic Barrier

**DOI:** 10.1101/667816

**Authors:** Z. E. Dell, M. Muthukumar

**Affiliations:** University of Massachusetts

## Abstract

Many fundamental biological processes involve moving macromolecules across membranes, through nanopores, in a process called translocation. Such motion is necessary for gene expression and regulation, tissue formation, and viral infection. Furthermore, in recent years nanopore technologies have been developed for single molecule detection of biological and synthetic macromolecules, which have been most notably employed in next generation DNA sequencing devices. Many successful theories have been established, which calculate the entropic barrier required to elongate a chain during translocation. However, these theories are at the level of the translocation coordinate (number of forward steps) and thus lack a clear connection to experiments and simulations. Furthermore, the proper diffusion coefficient for such a coordinate is unclear. In order to address these issues, we propose a center of mass (CM) theory for translocation. We start with the entropic barrier approach and show that the translocation coordinate is equivalent to the center of mass of the chain, providing a direct interpretation of previous theoretical studies. We thus recognize that the appropriate dynamics is given by CM diffusion, and calculate the appropriate diffusion constant (Rouse or Zimm) as the chain translocates. We illustrate our theoretical approach with a planar nanopore geometry and calculate some characteristic dynamical predictions. Our main result is the connection between the translocation coordinate and the chain CM, however, we also find that the translocation time is sped up by 1–2 orders of magnitude if hydrodynamic interactions are present. Our approach can be extended to include the details included in previous translocation theories. Most importantly this work provides a direct connection between theoretical approaches and experiments or simulations.

**SIGNIFICANCE:** Macromolecule motion through nanopores is critical for many biological processes, and has been recently employed for nucleic acid sequencing. Despite this, direct theoretical understandings of translocation are difficult to evaluate due to the introduction of the translocation coordinate. In this manuscript, we propose a theory for translocation written at the center of mass level of the polymer chain. This theoretical approach is more easily compared to experimental and simulation results, and additionally allows one to accurately account for hydrodynamic interactions on the macromolecule dynamics.

## INTRODUCTION

Many native biological processes are based on the manipulation of macromolecules through confined environments and the threading of macromolecules through nanopores, so called translocation (1–11). In eukaryotes, many fundamental processes in the cell require the transmission of nucleic acids or proteins across the nuclear membrane, between the cell nucleus and the cytoplasm. For example, after translation messenger RNA (mRNA) must move out of the nucleus into the cytoplasm, in order to be translated into proteins (2–4, 6). On the other hand, the proteins required for replication, transcription, and gene regulation have to enter the nucleus in order to interact with DNA and RNA. Such motion occurs through and is regulated by protein pores (5, 7–9). Similar to eukaryotic processes, many functions in prokaryotes and viruses require the motion of macromolecules across membranes, through small pores. For example, some viruses inject their genetic information into host cells in order to replicate (1, 10, 11). The knowledge and control of such processes, can lead to improved therapeutics and medicines. Thus, a thorough understanding of macromolecular motion during translocation through a nanopore is crucial to further biology, biophysics, and medicine.

In addition to fundamental biological processes, in recent years there has also been a significant increase in nanopore-based biotechnologies, which use protein or synthetic pores to manipulate and detect biological macromolecules (12–20). Most prominently this has been employed for DNA sequencing, which has led to the formation of several companies selling next generation sequencing devices (13, 18–20). Furthermore, adaptations of these devices to study RNA and protein structures is currently being explored (15, 17, 21). These biotechnologies in genomics and proteomics are an important aspect to advance personalized medicine.

In addition to the slew of biological applications, nanopores also have applications throughout synthetic materials and polymers (22–26). Borrowing ideas from biology, membranes and nanopores can be employed to direct bulk and selective transport of macromolecules through materials. Additionally, nanopore confinement can provide avenues of manipulating macromolecules for self-assembly and complexation (22, 23, 23, 24). From a fundamental polymer science prospective, it is important to understand the interplay between the entropic barriers required to confine macromolecules and the energetic driving forces due to chemical, electrical, and density gradients (25, 26). With a more complete understanding of these various contributions to translocation, one can develop better materials and biotechnologies.

Previous theoretical approaches to understand the translocation of a polymer chain can be categorized into two broad categories: (i) phenomenological or scaling-based theories (27–32) and (ii) microscopic or free energy-based theories (33–41). In many of the phenomenological theories, translocation proceeds due to tension propagation along the chain. Furthermore, polymer statistics are incorporated through the so-called tension blobs (27, 29–32). While this and other related approaches have some qualitative agreement with experiments, direct quantitative comparisons are difficult due to undetermined prefactors in scaling relations. Microscopic theories, on the other hand, formulate translocation in terms of Fokker–Planck dynamics, or drift–diffusion, where polymer effects are captured by the presence of an entropic barrier to translocation (26, 33–41). These theories are solved in various dynamical regimes (drift dominated, diffusion dominated) and for various chain models (flexible chains, rigid rods), leading to a vast array of predictions. However, most of these theories are developed at the level of the translocation coordinate, defined as the number of forward steps a polymer takes. Due to this formulation, two questions arise. First what is the interpretation of the translocation coordinate in terms of experiments or simulations? Namely, what actually constitutes a forward step? Second, what is the proper dynamical coefficient (diffusion constant) for the translocation coordinate? Previous work debates various answers to these questions, but mostly in an ad hoc manner (26, 32, 35). Furthermore, recent simulations indicate the importance of including proper hydrodynamic effects in order to predict the translocation behavior found in experiments (28, 42–44).

In this paper, we propose a theory for polymer translocation which resolves the two questions above. Our theory is based on mapping the previous microscopic, Fokker–Planck-based theories to the center of mass (CM) of the chain. To do so, we adopt a quasi-equilibrium assumption, which assumes that translocation proceeds slow enough that the chain can explore many configurations, and thus we calculate the CM of the chain for a given translocation coordinate. The quasi-equilibrium assumption is consistent with the previous formulation of the entropic barrier. When the theory is applied to a planar geometry, we find that the center of mass is a nearly linear function of the translocation coordinate. This indicates that the translocation coordinate can be directly interpreted as the center of mass, which is the primary result of this work. With the established mapping, the diffusion constant corresponding to translocation is no longer ambiguous, and must be the center of mass diffusion constant for the chain. We further, calculate the CM diffusion constant in the Rouse and Zimm models as the chain translocates and apply standard Langevin approaches to calculate the translocation time with and without driving fields. Several characteristic calculations are shown for planar geometry and indicate that by hydrodynamics speed up translocation.

The remainder of the paper is organized as follows. In the following section, we outline our theoretical approach in general and then apply it to study a planar nanopore geometry. This is followed by the results for the center of mass mapping and some characteristic translocation calculations. Finally, we conclude the paper with a summary and outlook to future work.

## THEORETICAL METHODS

In this section, we describe our theoretical approach for determining the center of mass (CM) dynamics of a polymer chain as it translocates through a nanopore. In the first subsection, we describe the model system and overview the general theoretical approach, the Fokker–Planck equation. The following three subsections describe the three components necessary to implement the Fokker–Planck approach. In the final subsection, we apply the technique to study a nanopore in an infinite planar geometry.

### Overview of the Center of Mass Fokker–Planck Approach

Consider a chain of *N* Kuhn segments (Kuhn length *l*_*k*_) that undergoes translocation through a geometric nanopore with diameter *d* ≈ *l*_*k*_ and length 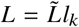, where 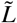 denotes the dimensionless pore length (Figure 1). We define the origin of our coordinate system at the center of the pore with the 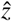 axis aligned along the pore axis.

**Figure 1:**
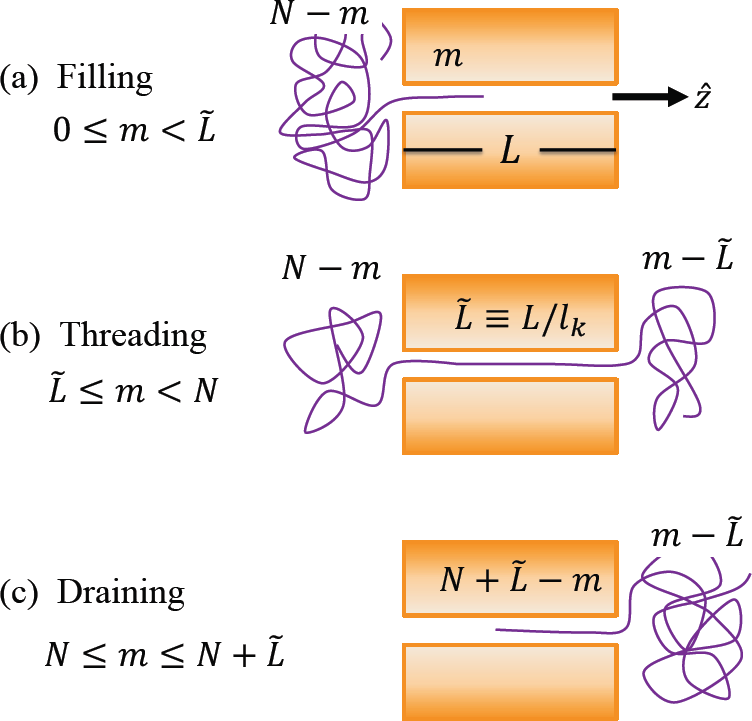
A schematic of a polymer chain (*N* Kuhn segments, each with Kuhn length *l*_*k*_) as it translocates through a nanopore (length *L*). There are three stages of translocation, characterized by the translocation coordinate *m*: (i) pore filling, (ii) chain threading, and (iii) pore draining.

Previous theories for translocation, are founded on calculating the free energy *F* (*m*) of the chain as it threads through the pore, where *m* is the translocation coordinate (26, 35). Theoretically, the translocation coordinate, 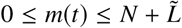, is the number of “forward Kuhn steps” the polymer has taken at time *t*, given *m*(0) = 0. To model the dynamics, a one dimensional Fokker–Planck equation for *m* (*t*) is established, with a reflecting boundary at *m* = 0 and an absorbing boundary at 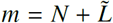 (45). The mean first passage time is then associated with the average translocation time (45, 46).

The most notable issue with the previous theoretical formulation for translocation is that there is no clean way to directly measure the translocation coordinate in experiments or simulations. Furthermore, this leads to an ambiguity in the choice for the effective diffusion constant of the translocation coordinate. Should it be related to the segmental motion of the segments inside the pore, or do the motions of the ends of the chain in the two chambers contribute to translocation dynamics? In the latter case, an understanding of chain dynamics on intermediate and center of mass length scales is necessary to fully describe translocation. In order to address these issues, in this work we develop a center of mass theory for translocation.

The starting point for our CM theory is the Fokker–Planck equation for the probability density 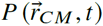 that the center of mass is located at position 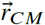 at time *t* (45):

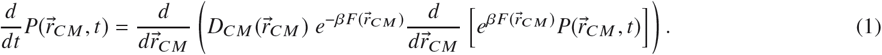

Here, *β* = 1/*k*_*B*_*T* is the inverse thermal energy, *F* is the free energy associated with the chain traversing the pore, and *D*_*CM*_ is the CM diffusion constant. The latter two quantities, in general, depend on the CM position as the chain translocates. Finally,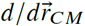 is the gradient operator for the CM coordinate.

When confined in the pore, by symmetry the average chain CM will fall along the pore axis 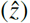, such that *x*_*CM*_ = 0 = *y*_*CM*_. In this case, Equation (1) reduces to a one dimensional equation:

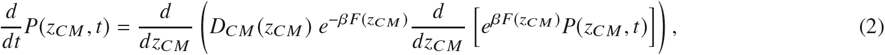

where *z*_*CM*_ = 0 when the center of mass is at the center of the pore; at the midpoint of translocation. The mean first passage time for the chain to fully translocate, move from *z*_*CM*_ =−*z*_0_ to +*z*_0_ (the initial and final positions of the chain, respectively), is thus given by:

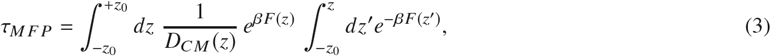

where we assume there is a reflecting boundary at −*z*_0_ and an absorbing boundary at +*z*_0_. The reflecting boundary prevents the chain from exiting into the initial chamber, since we are not interested in studying the capture of the chain. Thus, we associate τ_*M F P*_ with the mean translocation time.

We proceed by calculating the following three pieces: (i) the free energy as a function of the translocation coordinate *βF*(*m*), (ii) the center of mass diffusion constant as a function of the translocation coordinate *D*_*C M*_ (*m*), and (iii) a mapping between the center of mass and translocation coordinate *z*_*C M*_ ↔ *m*. In the following sections, we outline these three approaches, and then apply the results to study a planar geometry.

### Free Energy of a Translocating Chain

The free energy for a translocating chain will in general have three contributions (26, 33–35): (i) an entropic part associated with the portions of the chain outside the pore, (ii) an enthalpic part due to the pore–polymer interactions, and (iii) an enthalpic part due to driving forces (external fields, concentration gradients, etc.). By following previous theories (26, 33–35), the free energy can be calculated for various situations.

First, we consider a homopolymer translocating through a uniform pore, in the absence of any external driving forces. In this case, the free energy is divided into three sections of the translocation process: pore filling 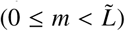, chain threading 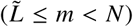, and pore draining 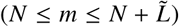, which are sketched in Figure 1. Thus, the free energy *F* is expressed as:

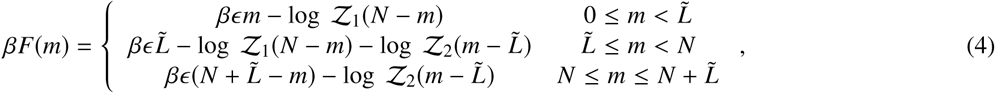

where ϵ is the interaction energy between a single Kuhn segment and the pore, which here is assumed to be constant throughout the pore. Furthermore, 𝒵_*i*_ is the partition function associated with a tethered chain in chamber *i* (where *i* = 1 and 2 denote the initial and final chambers, respectively). That is, log *Z* captures the entropy of the portions of the chain outside the pore and depends on the explicit geometry of the nanopore (see the discussion for planar geometry below).

In the presence of external driving, an explicit model of the driving force is necessary to properly model the free energy. For the commonly employed voltage-based driving of charged polymers, a simplified model is often adopted where the voltage in either chamber is taken as a constant and inside the pore there is a linear voltage ramp:

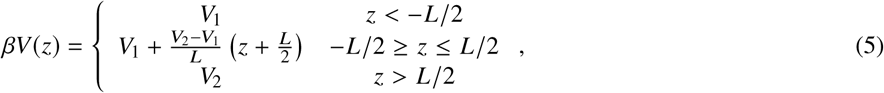

where *V*_1_ and *V*_2_ are the voltages in the two chambers, *z* < −*L/*2 and *z* > +*L/*2, respectively. By assuming each Kuhn segment has a charge *q*_*k*_ and that the chain inside the pore is rod-like, the free energy due to this voltage is easily expressed as a function of the translocation coordinate:

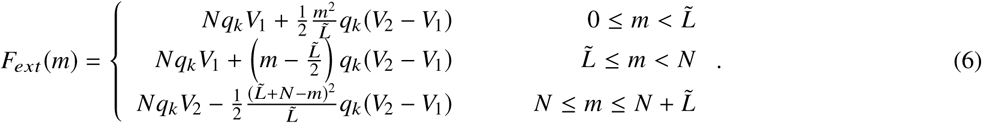

The total free energy under external driving is then obtained by adding Equations (4) and (6).

To completely specify the free energy, the entropic contributions must be calculated in terms of the translocation coordinate, by determining the partition function 𝒵_*i*_ (*m*) of a tethered chain with *m* Kuhn segments in the appropriate geometry. This can be done using the well-established Green’s function approach, which will be discussed in the context of planar geometries below.

### Center of Mass Diffusion Constant during Translocation

In addition to the free energy of the chain, we also need a model for the center of mass diffusion constant, *D*_*CM*_, as the chain translocates. We employ the Rouse–Zimm theory to calculate this (47–50). The CM diffusion constant of a chain is given by the Einstein relation *D*_*CM*_ = *k*_*B*_*T/ζ*_*CM*_, where *ζ*_*CM*_ is the center of mass friction on the chain. To calculate the friction, we first consider the Oseen tensor for two interacting segments (47, 50):

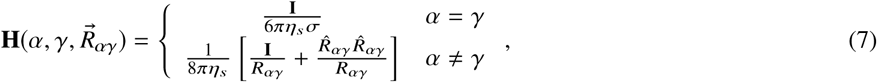

where α, γ ∈ [0, *N*] are the Kuhn segment labels for the two segments, 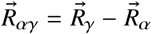 is the separation vector between the two segments, σ is the segment bead diameter, *η*_*s*_ is the solvent viscosity, and **I** is the identity tensor. Furthermore, we adopt the notation that 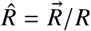 denotes a unit vector and *R* denotes the magnitude of a vector.

Before proceeding for a translocating chain, we first review the theoretical approach in the case of an isolated chain. We assume the system is isotropic and pre-average the Oseen tensor over all possible directions 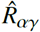. This results in 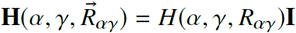, where the isotropic Oseen function is given by:

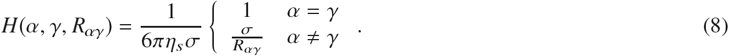

In the Rouse–Zimm theory (47, 50), we continue by pre-averaging the Oseen function over the chain segment distribution, which we denote by ⟨*H*⟩ and which depends only on |α γ|. From Equation (8), it is clear that ⟨1/*R*_αγ_⟩ is needed, which is usually evaluated via scaling arguments:

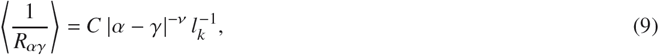

where *C* is a constant, and ν is the polymer size scaling exponent. In theta solvent ν = 1/2 and 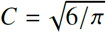, while in good solvent ν = 3/5 and the constant *C* is not easily determined (47–50).

Given the pre-averaged Oseen tensor, the center of mass friction on the chain is determined by (47, 50):

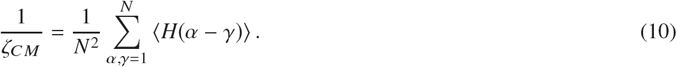

Combining Equations (8)–(10), approximating the double sum as a double integral, and taking the long chain limit *N* >> 1 yields:

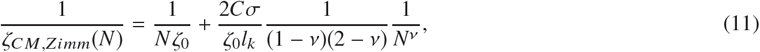

where *ζ*_0_ = 6*πη*_*s*_σ is the segmental friction constant. The first term in this expression is the Rouse friction coming from diagonal terms (no hydrodynamic interactions), while the second term is the Zimm contribution. In the long chain limit, the Zimm friction dominates since ν < 1, recovering *ζ*_*CM,Zimm*_ ~ *N*^ν^ as expected (47–50).

In order to apply this theoretical approach to the translocating chain, we adopt the following two assumptions: (i) the nanopore screens out hydrodynamic interactions, and (ii) the system effectively remains isotropic. Of these assumptions, the second is clearly problematic as the nanopore and any applied field will both introduce anisotropies. That being said, local hydrodynamic interactions will be isotropic and the hope is that this simple assumption will capture the leading order effects. In future work, a more complete anisotropic treatment can be developed using a tensorial approach.

Proceeding with the Rouse–Zimm analysis, the isotropic assumption implies that we can immediately reduce the Oseen tensor to the Oseen function as in Equation (8) and the CM friction is thus given by Equation (10). Given the assumption that hydrodynamics are screened by the nanopore, we can divide the chain into separate portions which internally have hydrodynamic interactions but do not interact with each other. We demonstrate this idea in Figure 2 for a general chain with two non-interacting portions. In this case, the Oseen function will take a block-diagonal form in terms of the bead labels α and γ. If, for generality, we assume there are *n* non-interacting chain portions (blocks), the CM friction reduces to:

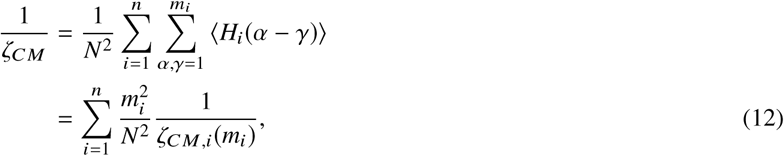

**Figure 2:**
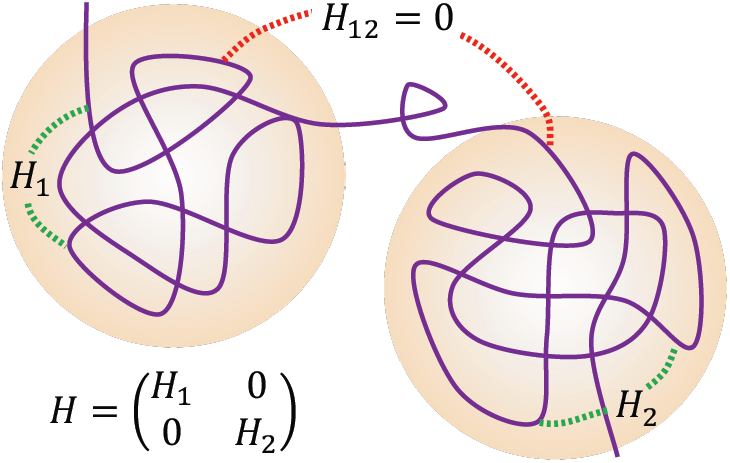
A schematic of two portions (orange spheres) of a polymer chain which do not hydrodynamically interact with each other. The hydrodynamics internal to each portion (green dotted lines) are characterized by a corresponding Oseen tensor (*H*_1_ and *H*_2_). However, the two portions do not interact corresponding to a zero-valued Oseen tensor (red dotted line, *H*_12_ = 0). Thus the full Oseen tensor for the chain, *H*, takes a block diagonal form.

where *m*_*i*_ is the total number of beads in chain block *i*, and *H*_*i*_ and *ζ*_*CM,i*_ are the corresponding Oseen function and center of mass friction for chain block *i*, respectively. There is an additional constraint that the total number of beads on the chain is fixed, 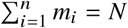. Equation (12) states that in order to determine the CM friction for the entire chain, we can calculate a weighted average of the CM frictions of the non-interacting portions of the chain. We note that this expression is general for any situation where hydrodynamic interactions between different parts of the chain are screened, resulting in a block-diagonal form of the Oseen function.

We proceed to apply Equation (12) to the translocating chain, which can be broken up into two or three portions depending on the stage of translocation (two for pore filling/draining and three for threading). For the portions of the chain outside the pore, we further assume that the chamber geometry and the pore do not affect the hydrodynamic interactions and thus, adopt the Rouse–Zimm model for their CM frictions (Equation (11)). For the portion of the chain inside the nanopore, given hydrodynamic interactions are screened, we model the CM friction as that of a rigid rod *ζ*_*C M,rod*_(*N*)= *N ζ*_0_. Combining these models with Equation (12) yields:

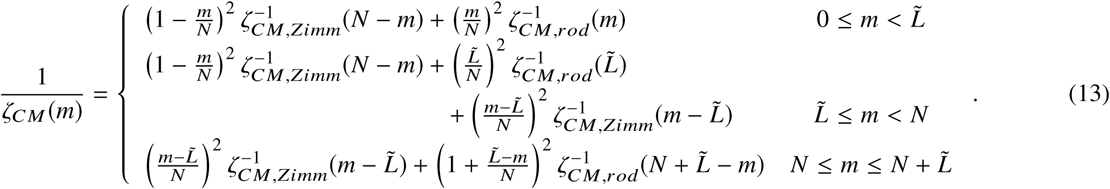

Equation (13), with the appropriate expressions for the rod and Zimm frictions, completes the theory for the center of mass diffusion constant, since *D*_*CM*_ = *k*_*B*_*T* /*ζ*_*CM*_.

### Relation between Center of Mass Position and the Translocation Coordinate

We now have expressions for the CM diffusion constant and the free energy of a chain as it translocates, in terms of the translocation coordinate *m*. In order to fully re-express the theory at the center of mass level, we need a mapping between the translocation coordinate and the center of mass position *m* ↔ *z*_*CM*_. We note that at the outset, it is not obvious that a one-to-one mapping is possible, however it seems physically reasonable. Furthermore, the results of our calculations demonstrate that such a mapping is indeed one-to-one (see the first subsection of the Results and Figure 4). This further implies that the translocation coordinate can be interpreted as analogous to the center of mass coordinate and provides a direct physical interpretation of the previous translocation theories.

**Figure 3:**
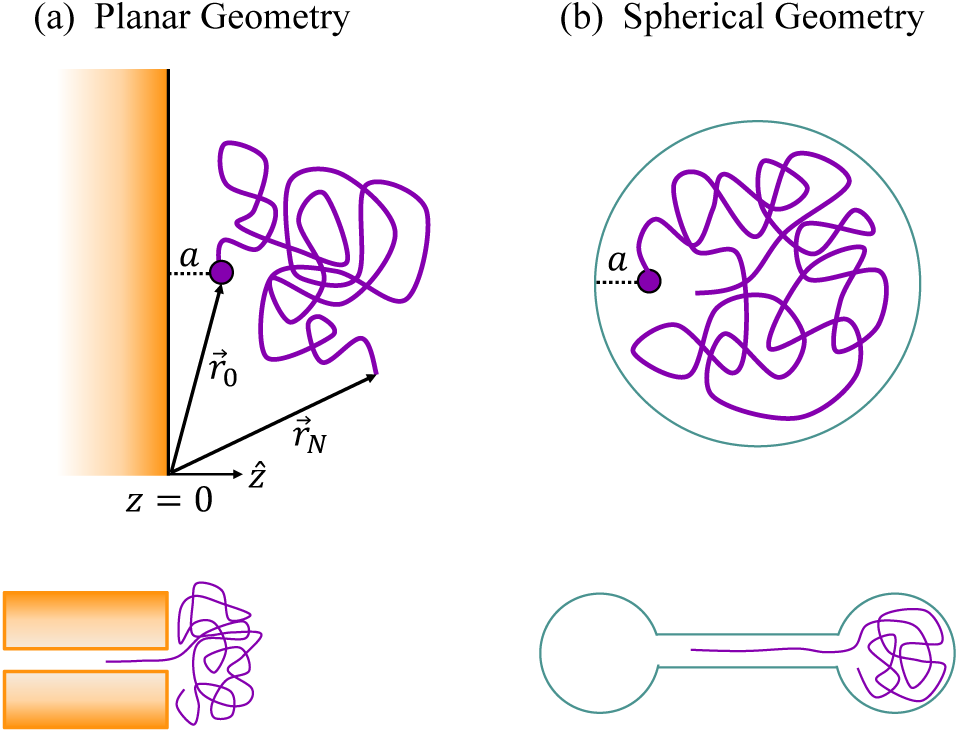
Two tethering geometries corresponding to typical nanopore configurations: (a) planar and (b) spherical. The chain ends are at positions 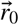 and 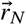, where one end is tethered a distance *a* away from the corresponding surface. For the planar geometry, we define the wall by *z* = 0.

**Figure 4:**
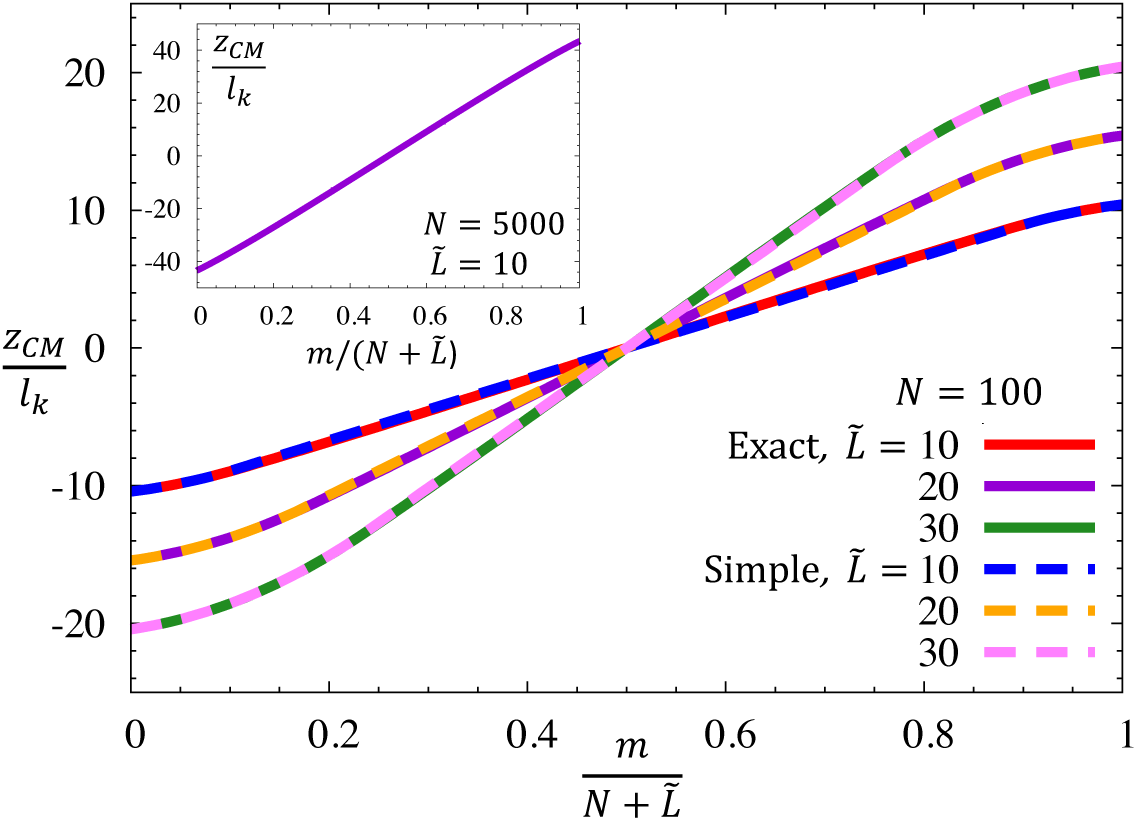
The center of mass of a polymer chain normalized by the Kuhn length *z*_*CM*_/*l*_*k*_ as a function of the translocation coordinate *m* normalized by its maximum value 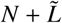 in a planar geometry. *z* = 0 corresponds to the center of the pore. (main) Results for a chain of *N* = 100 Kuhn segments and various dimensionless pore lengths 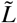. The solid curves show the exact results (Equation (26)) while the dashed curves show the results of our simplified model. (inset) Exact result for *N* = 5000 and 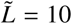.

To establish such a mapping, we calculate the center of mass of a chain after *m* forward steps of translocation. For a homopolymer, all beads have an identical mass and thus:

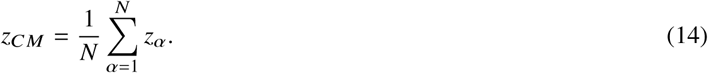

Dividing the chain into different portions in a manner analogous to the process we followed for the diffusion constant (although here the results do not depend on specific assumptions about chain interactions) yields:

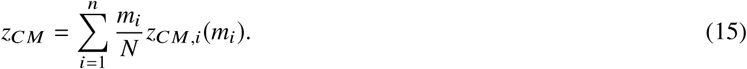

Thus, the center of mass is expressed as a weighted average of the center of masses of different portions of the chain.

In order to proceed, we must calculate the center of mass for two cases: (i) the parts of the chain that are outside of the pore; and (ii) the chain inside the pore. Given that many chain conformations can correspond to a single translocation coordinate *m*, we further adopt a quasi-equilibrium approximation; we assume that the chain will have ample time to explore many conformations before moving forward or backward. This assumption allows us to pre-average the center of mass coordinate over an equilibrium distribution at any given time step. While the validity of such quasi-equilibrium and pre-averaging is continuously debated, this assumption allows us to derive an effective Langevin equation for the CM coordinate, and is consistent with the average treatment used to determine the free energy.

To model the polymer inside the nanopore, we adopt a simple rod-like model, due to the narrow nature of the pore. Thus, the center of mass will simply be the center of the rod at any given instant, and can be determined in a given coordinate system. For the polymer outside of the pore, we must calculate the center of mass of a chain that is tethered near the pore opening, which depends on the chamber geometry. We also ignore the opening of the pore as a second order correction.

We outline the approach for calculating the center of mass in a generic chamber here and apply the approach to a specific geometry in the following section. The center of mass position can be determined in terms of the segment density 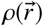, via:

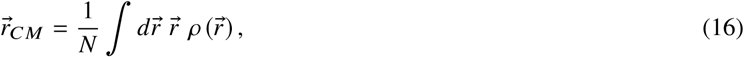

which is a natural generalization of Equation (14). Furthermore, for a tethered chain (here effectively tethered by the mouth of the pore) the segment density is determined by the Green’s function, *G* via:

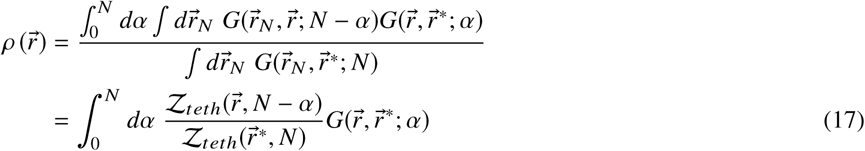

where a continuum, space curve, approximation for bead label α is made. 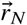 and 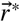 represent the position of the free end and the tether, respectively. The Green’s function 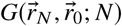 is the probability density of having a chain of length *N* that has one endat position 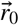 and the other end at position 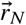; and the tethering partition function is given by 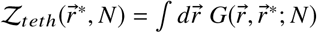, which is symmetric in integrating over either endpoint.

With the above approach, the centers of mass in the two chambers can be calculated. Furthermore, we adopt the coordinate system with origin at the center of the pore. Thus the translocation center of mass is easily determined via:

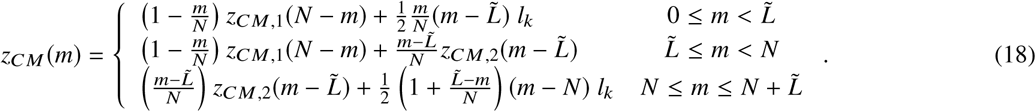

The center of mass in the two chambers *z*_*CM,i*_ will be calculated for an example geometry in the following subsection.

### Theory for a Planar Geometry

We now apply the theory outlined in the previous subsections to a specific geometry. There are two commonly employed geometries for translocation (Figure 3) (35, 37). The first case is a planar geometry (Figure 3a), which models a nanopore embedded in a flat lipid bilayer or a solid state nanopore. A second common case is the spherical geometry (Figure 3b), which models macromolecule expulsion from a virus or drug capsid. In this paper we focus only on the planar case, but note that the generalization to the spherical geometry is straightforward.

The free energy and the chain center of mass are both determined by the Green’s function for a tethered chain. In order to calculate the Green’s function for the planar geometry, we adopt a Gaussian approximation for the chain, since it allows for analytic tractability. In future work, we will explore the consequences of this approximation and consider other models such as excluded volume chains. The Green’s function of an isolated Gaussian chain (not near a wall) is (26, 47, 51):

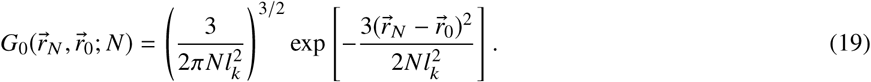

To model the wall, we adopt an infinite planar geometry defined by the *z* = 0 plane, as shown in Figure 3. The normal to the wall is 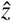. Thus, the Green’s function must satisfy the boundary condition:

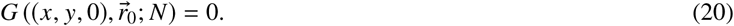

We assume that by construction 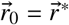; the tether is not on the wall.

In order to satisfy the condition in Equation (20), the method of images can be applied (52, 53). We imagine the wall does not exist and consider two chains both with an end at 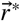. The first chain is the one of interest, with the other end at 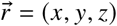, while the second chain is an image chain, which terminates at 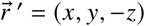 (reflected across the wall). By considering these chains, the Green’s function in the presence of the wall is:

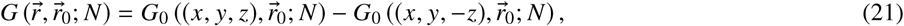

where *G*_0_ is the isolated chain (no wall) Green’s function. Equation (21) clearly satisfies both the diffusion equation and the boundary condition, and by uniqueness is the solution to the question at hand.

From the Green’s function, we calculate the tethering partition function via:

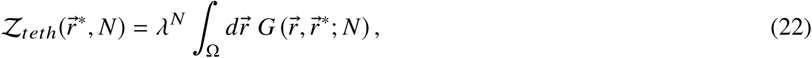

where the volume Ω is the half-space that the chain is in, *z* > 0. Additionally, *λ* is a factor which relates the Green’s function to the number of conformations of the chain. Typically, *λ* is thought of either as the coordination number in a lattice-based theory or as the fugacity *λ* = exp(*βµ*), where *µ* is the chemical potential (26, 51). From Equations (19) and (21), we can directly evaluate Equation (22), which yields:

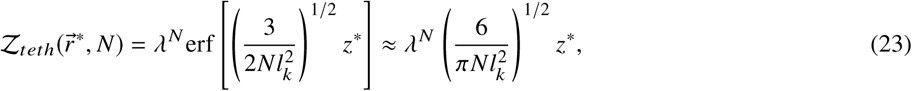

where erf(…)is the error function, and the approximate equality is for a tether close to the wall, for *z*\*<< *N*^1/2^*l*_*k*_. With this partition sum, the free energy is easily calculated.

The center of mass calculation for the planar geometry involves employing Equations (21) and (23) in Equations (16) and (17). This calculation is described in the Appendix and here we outline the main results. By symmetry, and additionally verified by integration in our model, *x*_*CM*_ = *x*\* and *y*_*CM*_ = *y*\*; the center of mass is aligned with the tethering site in the 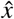 and 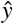 directions. In the 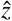 direction, the calculation is complicated by the presence of the wall.

An analytic solution is not possible for a general tethering position. However, for translocation, tethering is done very close to the wall and thus the integrals can be expanded about the *z*\* << *N*^1/2^*l*_*k*_ limit. With this approximation, the integrals are analytically possible (see the Appendix) and yield:

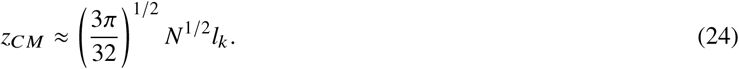

Hence, when the tether is very close to the wall, the center of mass scales as the radius of gyration, *z*_*CM*_ ~ *N*^1/2^ ∼ *R*_*g*_. This scaling is intuitive and a general result, however determining *R*_*g*_ near a wall is often complex for non-Gaussian models.

Summarizing the main results for translocation of a chain through a pore imbedded in a plane. The free energy in the absence of an applied field, as a function of the translocation coordinate, follows from Equations (4) and (23):

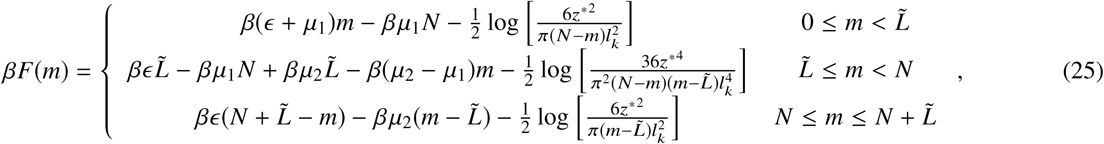

where *µ*_1_ and *µ*_2_ are the chemical potentials for the Kuhn segments in the initial and final chambers, respectively.

Likewise, the center of mass follows from Equations (18) and (24), where the origin is taken at the center of the pore:

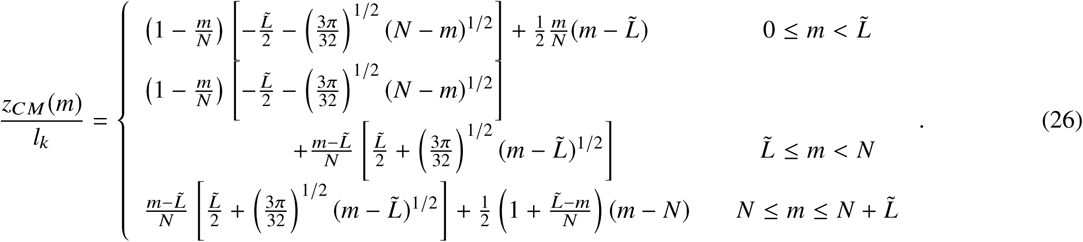

Given the free energy in Equation (25), the CM diffusion constant described in the section above, and the inverse of Equation (26) (the translocation coordinate corresponding to a given center of mass), the CM dynamics theory described at the beginning of the methods is closed.

## RESULTS AND DISCUSSION FOR PLANAR GEOMETRY

In this section, we present some results for the translocation of a chain through a planar geometry. We begin by showing the center of mass in terms of the translocation coordinate and then develop a simple model for inverting this relation. Then we present some of the dynamical results.

### Mapping between the Center of Map and Translocation Coordinate

Before studying the translocation dynamics, we must establish the mapping between the translocation coordinate *m* and the center of mass position *z*_*CM*_. As previously stated, there is no guarantee that under our assumptions such a mapping will be one-to-one, however we will see this is indeed true.

Figure 4 shows the center of mass *z*_*CM*_ as a function of the translocation coordinate *m* for various chain lengths *N* and pore lengths *L*. In these results, all lengths are normalized by the Kuhn length *l*_*k*_ and the translocation coordinate is normalized by its maximum value 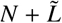, such that it falls in the range [0, 1]. The main frame shows the results for *N* = 100 and where the solid lines (red, purple, and green) indicate the exact results, using Equation (26), for various pore lengths (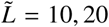 and 30, respectively). The inset shows the exact results for *N* = 5000 and 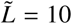. In all cases, the center of mass monotonically increases as translocation proceeds and is symmetric about 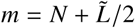 where *z*_*CM*_ = 0. For 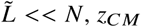 is nearly linear in *m* and for all other cases, during the threading stage this linearity holds.

For the translocation theory, we must invert this mapping. No direct analytic inverse of Equation (26) exists, however numerical inversion is possible. Rather than doing this, we exploit the nearly linear nature of the exact results in the threading regime and propose the following simplified model for the center of mass. In the threading state, *m* << *N* and thus we Taylor expand Equation (26) about *m* = 0, keeping out to second order in *m*, since the exact results are clearly nonlinear. Likewise, for the draining regime, we expand about 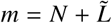. For the threading regime, we adopt a linear function which is uniquely determined by ensuring continuity between the threading result and the draining result. This simplified model for the center of mass is then:

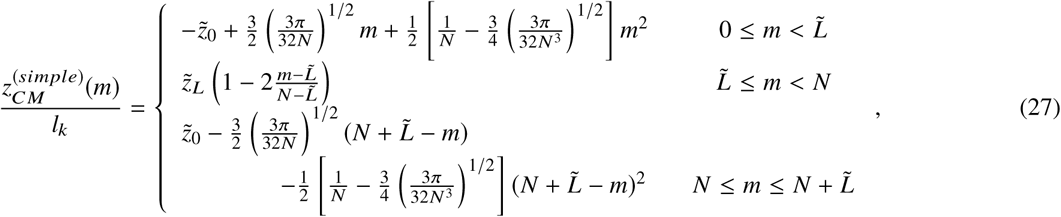

Where 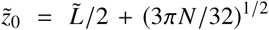 is the starting position of the center of mass, and 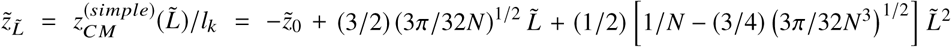 is the value of the simplified center of mass at the end of the filling regime. Note that the simplified model satisfies the symmetry about 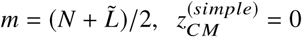, by construction.

To confirm that this simplified model accurately captures the behavior of the center of mass, we show the results of the simplified model as dashed lines in the main frame of Figure 4. We note that there is a very weak deviation from true linear behavior in the threading regime, however in general the simplified model is within 5% of the exact model. Thus in all that follows, we only use the simplified model for the center of mass.

Given Equation (27) is quadratic at highest order, a direct analytic inversion is possible, yielding:

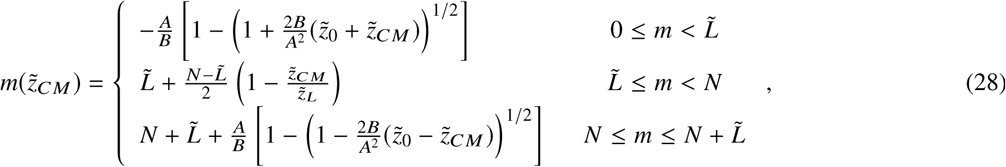

where 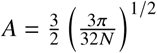 and 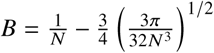. To derive the inversion given in Equation (28), we choose the quadratic roots such that 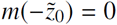 and 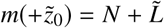.

Figure 4 in conjunction with Equations (27) and (28) provide the primary conclusion of this paper. While this result does not make any dynamical predictions for translocation, it provides the basis for a fundamental interpretation of the previous translocation theories. The previous theories are typically constructed in terms of the translocation coordinate *m*, which expresses the number of forward Kuhn steps the polymer takes as it translocates. While theoretically well-defined, this quantity is difficult to interpret in an experimental or simulation framework. Furthermore, the dynamical coefficient (effective diffusion constant) for this variable is somewhat ambiguous. Given the above results, we have now provided direct evidence that the translocation variable *m* is equivalent to the center of mass of the chain *z*_*CM*_ provided we adopt the quasi-equilibrium approximation. Even stronger than this, we have shown that in the limit that the chain length is much larger than the pore length,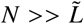, this equivalence is very close to linear.

This result allows us to directly connect the theory to experiments and simulations. Further, we can now predict translocation dynamics with the proper dynamical coefficient, the center of mass diffusion constant. The results for the center of mass translocation dynamics are described in the following subsection.

### Dynamic Properties in the Center of Mass Theory

In this section, we present some characteristic results for the dynamics of a translocating chain. We first focus on the free energy and CM diffusion constant, before making some predictions of the translocation time. These results are in no means a comprehensive study, but rather some characteristic examples of how the theory can be applied to a given translocation setup.

In all of the following results, we take the chemical potentials to be *µ*_1_ = *µ*_2_ = 0.01*k*_*B*_*T* and consider a neutral pore *β*∈ = 0,with a length of *L* = 10*l*_*k*_. Furthermore, we take the charge of a Kuhn segment to be *q*_*k*_ = −*e*, where *e* is the elementary charge, and the voltage difference to be *V*_2_ − *V*_1_ = *V*. Note, for the free energy, we consider the initial state of the chain as the reference Δ *F*(*z*) = *F*(*z*) − *F*(−*z*_0_), and thus the results depend only on the voltage difference *V*.

Figure 5 shows the translocation free energy profiles *β*Δ *F* as a function of the center of mass position *z*_*CM*_, for chain lengths *N* = 100, 200, 500, and 1000. The center of mass coordinate falls in the range [−*z*_0_, +*z*_0_], where 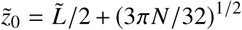. Since *z*_0_ ~ *N*^1/2^ for small pores lengths, we have normalized the center of mass coordinate by *z*_0_ so that the free energies can be directly compared. Thus, a value of −1 indicates the start of translocation and a value of +1 indicates the end of translocation.

**Figure 5:**
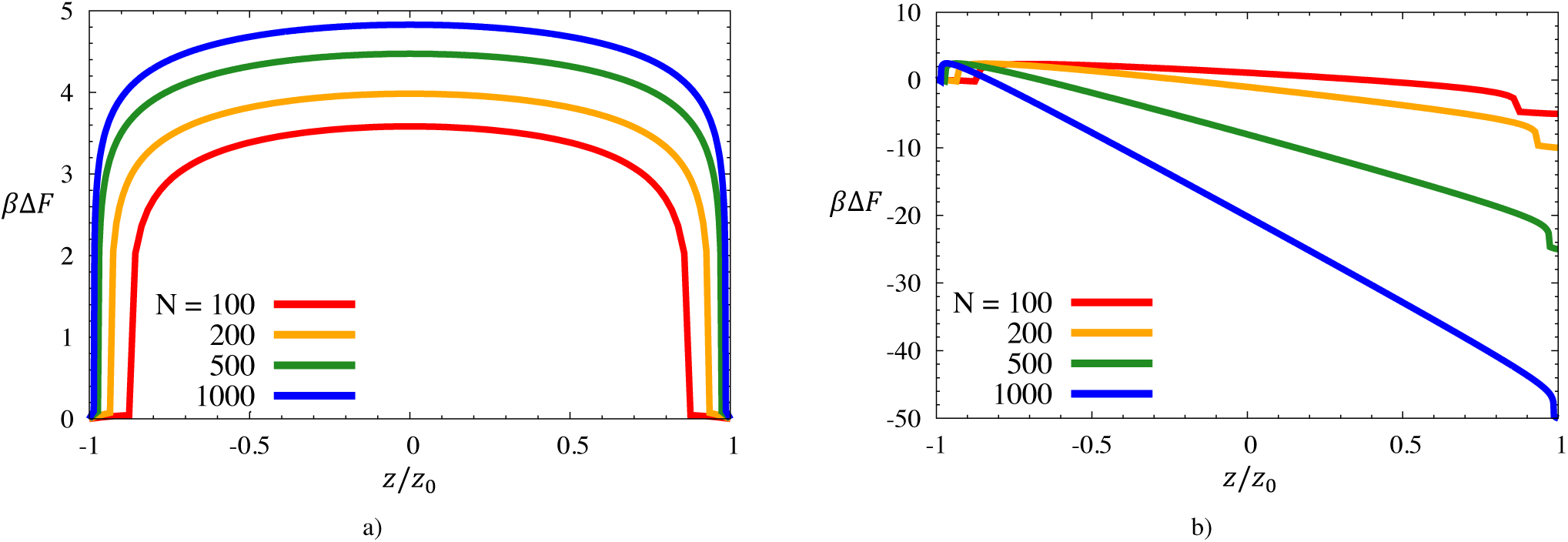
Translocation free energy, expressed in units of thermal energy *β* = 1/*k*_*B*_*T* and relative to the initial free energyΔ *F* = *F*(*z*)− *F* (–*z*_0_), as a function of the center of mass position, expressed relative to its initial distance from the center of the pore *z*_0_. Various chain lengths *N*, and two applied voltages are shown: (a) no applied voltage *V* = 0 (chain length increases from bottom curve to top curve), and (b) a high applied voltage *V* = 0.05 *k*_*B*_*T /e* (chain length increases from top curve to bottom curve).

We first consider the case when *V* = 0 (Figure 5a), which since *µ*_1_ = *µ*_2_ corresponds to no external driving force. In this case, the free energy is completely determined by the chain entropy. As the pore fills (most evident for *N* = 100, red curve), there is a weak increase in the free energy due to the entropy lost by restricting the first ten segments inside the pore. Once the pore is filled there is a dramatic increase in the free energy corresponding to the entropy lost when the chain is divided into two parts (those in the initial and final chambers). By symmetry, the maximum free energy is found when the chain is at the middle of the translocation process *z*_*CM*_ = 0. Here the chain is divided into two equal smaller chains (apart from the segments inside the pore), and Δ *F*(0) is the net barrier for translocation. The translocation barrier increases with chain length, from *F*_*B*_ ≈ 3.6*k*_*B*_*T* for *N* = 100 to *F*_*B*_ ≈ 4.8*k*_*B*_*T* for *N* = 1000, Figure 5a. While the barrier only weakly increases, this can have a dramatic effect on the relaxation time, since τ ~ exp(*βF*_*B*_). Finally, we point out that the biggest bottleneck to translocation is in the initial threading regime, *z*_*CM*_ ≲ −0.75*z*_0_. Here the force driving the chain back into the initial cavity (given by the slope of the free energy) is the largest. Likewise, after the chain has surpassed a certain threading threshold, *z*_*CM*_ ≲ +0.75*z*_0_, it will be quickly extracted from the pore due to strong entropic gains in exiting. For all curves in Figure 5, there is a dramatic change in behavior as the chain drains from the pore (*z* ≈ *z*_0_); the free energy only weakly varies in this regime. Additionally, as the chain length increases, the draining regime becomes negligible compared to the overall translocation process.

When a voltage of *V* = 0.05 *k*_*B*_*T/e* is applied across the pore (Figure 5b), the free energies shown in Figure 5a are further modified by the electrostatic contributions (Equation (6)). In this case, the driving forces due to the voltage dominate the free energy profiles. Note that as the chain length is increased, the effect of the voltage increases since there are more charges present on the chain; the overall free energy change is *F*(*z*_0_)−*F* (−*z*_0_)= −*NeV*. We point out that despite the presence of a driving field, in all cases there remains a roughly 3*k*_*B*_*T* barrier for the initial threading of the chain. While this barrier will ultimately affect the translocation process, once surpassed the chain will quickly translocate due to the driving field.

As is evident from Equation (3), the translocation time depends not only on the free energy, but also on the center of mass diffusion constant as the chain translocates. Previous theories for translocation were either based at the segmental level, with constant segmental diffusion constant *D*_0_, or equivalently at the center of mass level with only Rouse dynamics, where *D*_*CM*_ = *D*_0_/*N* regardless of the stage of translocation. In our proposed CM theory, we include the possibility of hydrodynamics outside the nanopore and thus have a Rouse–Zimm model where the diffusivity depends on the center of mass position. Figure 6 shows the results of our model (Equation (13) with the CM mapping) for various chain lengths in good solvent (taking the Zimm prefactor *C* = 1 and exponent ν = 3/5). In Figure 6, the center of mass diffusivity is normalized by the bulk Zimm diffusion constant *D*_*Zimm*_ = *k*_*B*_*T/ζ*_*CM,Zimm*_ *N*, where *ζ*_*CM,Zimm*_ *N* follows from Equation (11). Thus, at the beginning and end of translocation *D*_*CM*_ = *D*_*Zimm*_. As the chain translocates, its diffusion slows down until the midpoint of translocation, after which it speeds up. This effect is a direct consequence of the fact that hydrodynamic speed up is more prominent for longer chains. To demonstrate this further, consider the Zimm friction at the midpoint of translocation *ζ*_*CM*_ *(z*_*CM*_ = 0) ~ 2 * (*N/*2)^3^ 5 = 2^2/5^ *ζ*_*CM*_(*z*_0_), which is clearly larger than the full chain friction. Finally, note that there is a weak chain length dependence, especially for shorter chains, leading to slower diffusion. This is due a direct consequence of the fact that Rouse-like behavior becomes more prominent for short chains in Equation (11).

**Figure 6:**
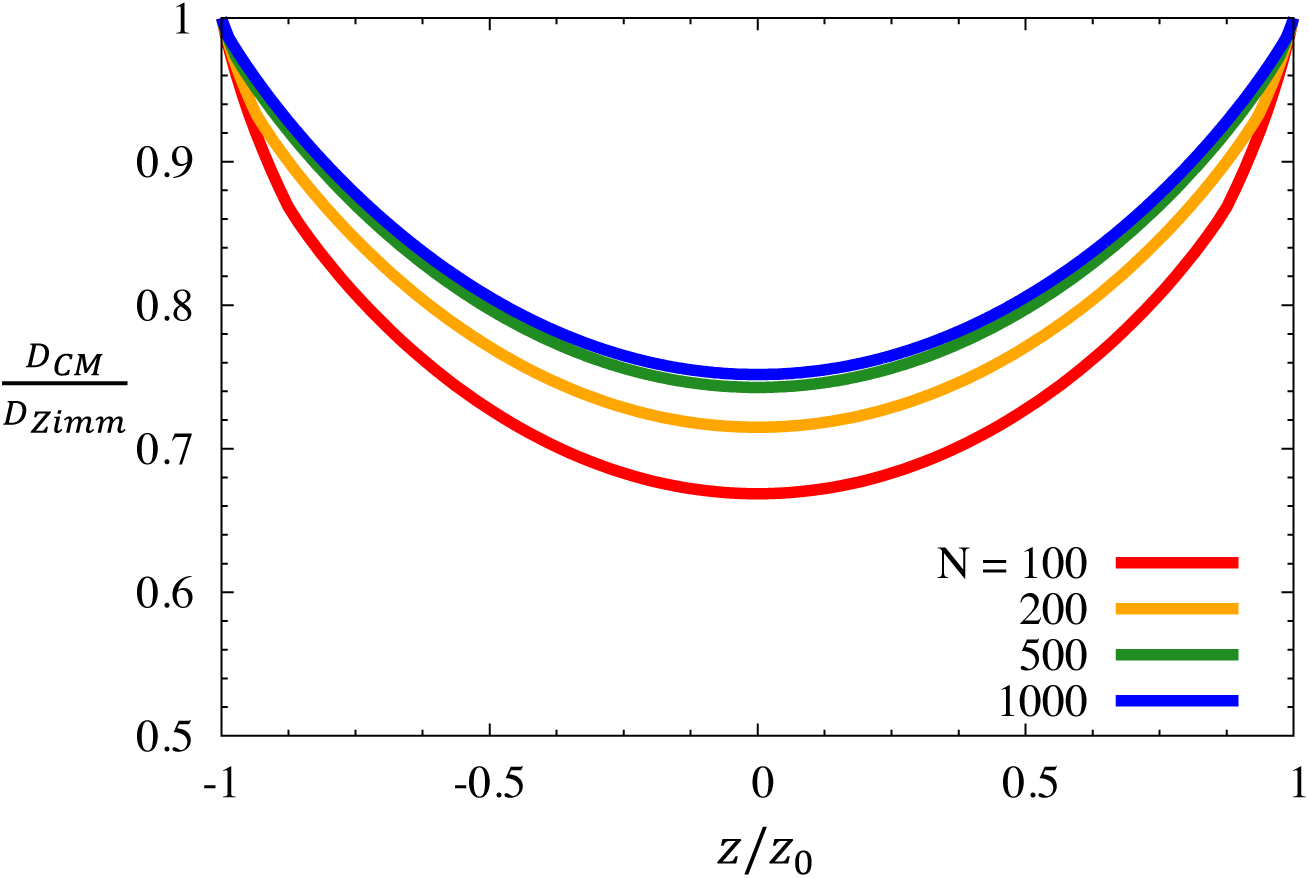
The center of mass diffusion constant *D*_*CM*_ for a Rouse–Zimm chain as it translocates, relative to the bulk Zimm diffusion constant for the same chain *D*_*Zimm*_ ~ *N*^*v*^, in good solvent (prefactor *C* = 1 and exponent ν = 3 5). Chain length increases from bottom curve to top curve.

Given the free energy and the CM diffusion constant, we can predict translocation times under various conditions from Equation (3). Figure 7 shows some characteristic results for the translocation time, relative to the segmental relaxation time 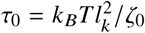. Figure 7a shows the results for various voltages as a function of the chain length *N*. The solid curves show the results including hydrodynamics using the modeled proposed here (Zimm label), while the dashed curves show the results without hydrodynamics using a simple Rouse diffusivity *D*_*CM*_ = *k*_*B*_*T/N ζ*_0_ for all *z*_*CM*_. Note that the Rouse model results are equivalent to previous theories. The red, top curves in both cases show the results for no driving field *V* = 0, and as the voltage is increased the translocation time decreases as expected. The chain length dependence for the Rouse and Zimm models are qualitatively similar, however, in the Zimm model the translocation time is significantly faster (1–2 orders of magnitude) due to hydrodynamic speed up. Additionally, we point out for the largest applied field under the Zimm model, the translocation time is nearly constant in *N*. This nearly constant translocation time is due to the complex interplay of the driving field, the entropic barrier, and hydrodynamic speed up for long chains.

**Figure 7:**
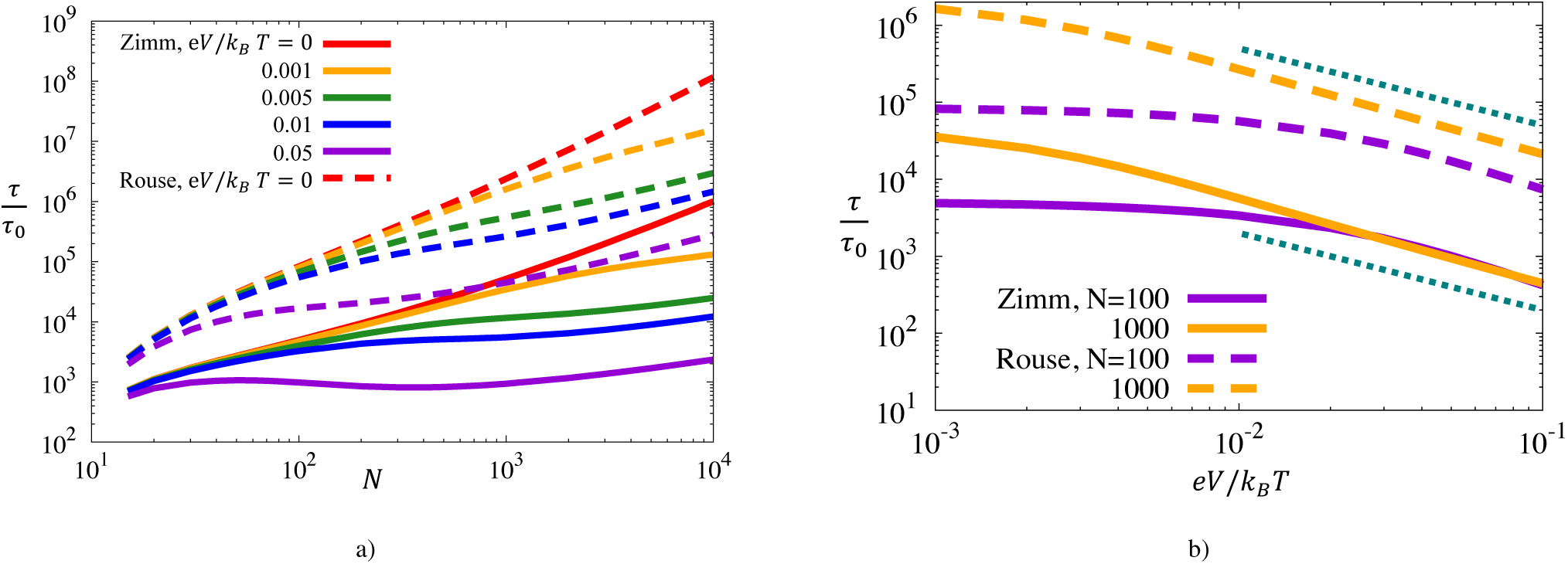
The mean first passage time (translocation time), expressed in units of segmental time 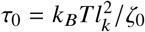 as a function of (a) chain length *N*, and (b) applied voltage *eV/k*_*B*_*T*. The dashed curves correspond to a Rouse chain in all three regions (the two chambers and the pore) such that *D*_*CM*_(*z*) ~ 1/*N* is constant. This model is equivalent to the previous translocation theories scaled by *N*. The solid curves assume the Rouse–Zimm model proposed here. In frame (a), the voltage increases from the top curve to the bottom curve for both models. In frame (b), the top curves are for *N* = 1000 in both models.

Finally, we analyze the translocation time as a function of the driving voltage *V* (Figure 7b) for *N* = 100 (purple curves) and *N* = 1000 (orange curves). As the voltage is increased, there is a clear decrease in the translocation time due to the stronger driving forces. In all cases, for large driving field this decrease is characterized by an inverse relationship τ ~ *V* ^−1^, which is shown by the dotted teal lines in the figure. From these characteristic results, it is clear that the translocation behavior of a chain in various situations can be described with our theoretical approach. Understanding these results is crucial for controlling translocation, which will ultimately lead to a more complete picture of biological processes and better development of biotechnologies.

## CONCLUSIONS

In this work, we have generalized the previous theory for translocation to the center of mass level by combining translocation theory with the Rouse–Zimm model for CM diffusion and by developing a mapping between the translocation coordinate *m* and the center of mass *z*_*CM*_. While the theoretical approach is rather general, here we applied it to the case of a planar nanopore in order to explicitly illustrate the approach. The primary result of this paper is that there is a one-to-one mapping between the translocation coordinate and the center of mass. Furthermore, when the chain contour length is large relative to the pore length *Nl*_*k*_ >> *L*, this mapping is approximately linear. While this result seems obvious, it actually has profound physical meaning; this linear mapping implies that the translocation coordinate, used in many theoretical approaches, directly captures center of mass behavior of the chain. Additionally, this provides a clear connection between translocation experiments and many theoretical descriptions.

Our theoretical approach is also able to properly incorporate hydrodynamic interactions, and several characteristic results for the translocation behavior were analyzed. In general, hydrodynamics sped up the translocation process by 1–2 orders of magnitude, and under certain situations even led to translocation times that were nearly independent of the chain length. These initial results demonstrate the importance of properly accounting for hydrodynamics when necessary.

In the future, this approach can be employed to study alternative geometries, as has been done with more classical translocation theories (35, 37). Furthermore, a more complete model of the center of mass position, which properly accounts for excluded volume interactions, can be included. This work provides the fundamental theoretical foundations for studying many other complex situations, such as those involving random or block co-polymer translocation (relevant to proteins) or center of mass polymer motion through porous environments. Additionally, this approach can be used to interpret experimental data related to translocation. With this better understanding of translocation from a theoretical perspective, one can design better methods for controlling polymer motion, develop better biotechnologies for nucleic acid and protein sequencing, and cultivate a better understanding of many biological processes.

## APPENDIX CENTER OF MASS CALCULATION FOR PLANAR GEOMETRY

For the center of mass, we need to first calculate the density by employing Equations (21) and (23) in Equation (17):

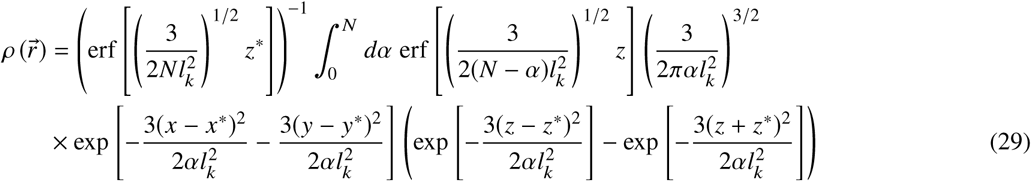

A general analytic result for the density is not possible, but can be easily numerically calculated.

Despite the lack of an analytic expression for the density, by using Equation (29) in Equation (16), we can solve for the center of mass:

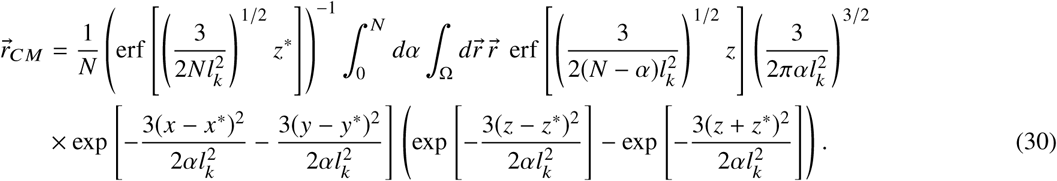

For the 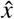 and 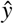 components, the integrals can be explicitly evaluated to yield *x*_*CM*_ = *x*\* and *y*_*CM*_ = *y*\* as expected by symmetry. For the 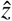 direction, the result is complicated due to the presence of the wall.

Again a general analytic expression is not possible, however if we assume the tether is very close to the wall, as relevant to translocation, then we can expand about *z*\* << α^1/2^*l*_*k*_ << *N*^1^ /2*l*_*k*_. Under this approximation, the general expression in Equation (30) reduces to:

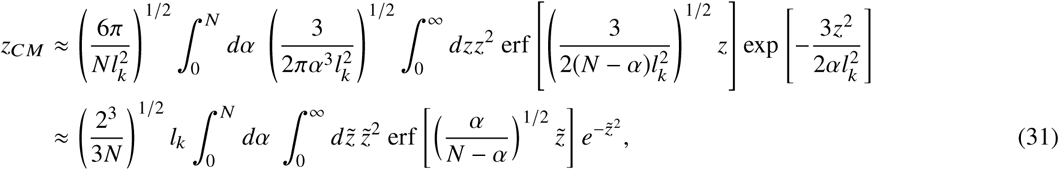

where the second line employs the coordinate transform 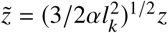. Performing the 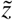 integral results in:

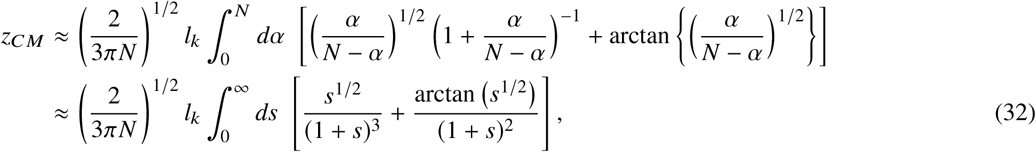

where the second line follows from *s* = α/(*N* − α). The resulting integrals are easily evaluated yielding:

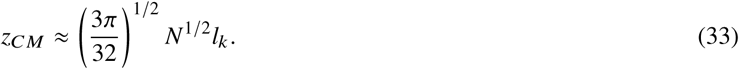

This is the result we stated in Equation (24) of the main text.

## AUTHOR CONTRIBUTIONS

ZED developed the theory, performed the calculations, analyzed the data, and wrote the manuscript. MM initiated the study, oversaw the theoretical development and analysis, and edited the manuscript.

## ACKNOWLEDGMENTS

The authors of this paper acknowledge Sabin Adhikari for useful discussions. This work was supported by the National Institutes of Health (grant no. R01HG002776-15), the National Science Foundation (grant no. DMR-1713696), and the Air Force Office of Scientific Research (grant no. FA9550-17-1-0160).

## REFERENCES

1. Alberts, B., A. Johnson, J. Lewis, M. Raff, K. Roberts, and P. Walter, 2007. Molecular Biology of the Cell. Garland Science, New York, NY, USA, 5th edition.

2. Li, Y., S. L. Junod, A. Ruba, J. M. Kelich, and W. Yang, 2018. Nuclear export of mRNA molecules studied by SPEED microscopy. Methods 153:46.

3. Tudek, A., M. Schmid, and T. H. Jensen, 2019. Escaping nuclear decay: the significance of mRNA export for gene expression. Current Genetics 65:473.

4. Harries, L. W., 2019. RNA biology provides new therapeutic targets for human disease. Frontiers in Genetics 10:205.

5. Glukhova, A., E. Nabirochkina, and D. Kopytova, 2018. mRNP Transport in Eukaryotes. mRNP Export from the Nucleus. Molecular Genetics, Microbiology and Virology 33:182.

6. Neriec, N., and P. Percipalle, 2018. Sorting mRNA molecules for cytoplasmic transport and localization. Frontiers in Genetics 9:510.

7. Hoelz, A., E. W. Debler, and G. Blobel, 2011. The structure of the nuclear pore complex. Annual Review of Biochemistry 80:613.

8. Beck, M., and E. Hurt, 2017. The nuclear pore complex: understanding its function through structural insight. Nature Reviews Molecular Cell Biology 18:73.

9. Eftekharzadeh, B., J. G. Daigle, L. E. Kapinos, A. Coyne, J. Schiantarelli, Y. Carlomagno, C. Cook, S. J. Miller, S. Dujardin, A. S. Amaral, et al., 2018. Tau protein disrupts nucleocytoplasmic transport in Alzheimer’s disease. Neuron 99:925.

10. Allemand, J.-F., B. Maier, and D. E. Smith, 2012. Molecular motors for DNA translocation in prokaryotes. Current Opinion in Biotechnology 23:503.

11. Nummela, J., and I. Andricioaei, 2008. Energy Landscape for DNA Rotation and Sliding through a Phage Portal. Biophysical Journal: Biophysical Letters 96:L29.

12. Kasianowicz, J. J., E. Brandin, D. Branton, and D. W. Deamer, 1996. Characterization of individual polynucleotide molecules using a membrane channel. Proceedings of the National Academy of Sciences 93:13770.

13. Meller, A., L. Nivon, E. Brandin, J. Golovchenko, and D. Branton, 2000. Rapid nanopore discrimination between single polynucleotide molecules. Proceedings of the National Academy of Sciences 97:1079.

14. Akeson, M., D. Branton, J. J. Kasianowicz, E. Brandin, and D. W. Deamer, 1999. Microsecond time-scale discrimination among polycytidylic acid, polyadenylic acid, and polyuridylic acid as homopolymers or as segments within single RNA molecules. Biophysical Journal 77:3227.

15. Butler, T. Z., J. H. Gundlach, and M. A. Troll, 2006. Determination of RNA orientation during translocation through a biological nanopore. Biophysical Journal 90:190.

16. Hartel, A. J., S. Shekar, P. Ong, I. Schroeder, G. Thiel, and K. L. Shepard, 2019. High bandwidth approaches in nanopore and ion channel recordings–A tutorial review. Analytica Chimica Acta 1061:13.

17. Nehra, A., S. Ahlawat, and K. P. Singh, 2019. A biosensing Expedition of Nanopore: A review. Sensors and Actuators B: Chemical 284:595.

18. Buermans, H., and J. Den Dunnen, 2014. Next generation sequencing technology: advances and applications. Biochimica et Biophysica Acta 1842:1932.

19. Haque, F., J. Li, H.-C. Wu, X.-J. Liang, and P. Guo, 2013. Solid-state and biological nanopore for real-time sensing of single chemical and sequencing of DNA. Nano Today 8:56.

20. Wanunu, M., 2012. Nanopores: A journey towards DNA sequencing. Physics of Life Reviews 9:125.

21. Wang, H., J. Ettedgui, J. Forstater, J. W. Robertson, J. E. Reiner, H. Zhang, S. Chen, and J. J. Kasianowicz, 2018. Determining the physical properties of molecules with nanometer-scale pores. ACS Sensors 3:251.

22. Howorka, S., and Z. Siwy, 2009. Nanopores: generation, engineering, and single-molecule applications. In Handbook of Single-Molecule Biophysics, Springer, 293.

23. Murphy, R. J., and M. Muthukumar, 2007. Threading synthetic polyelectrolytes through protein pores. Journal of Chemical Physics 126:051101.

24. Nakane, J., M. Akeson, and A. Marziali, 2002. Evaluation of nanopores as candidates for electronic analyte detection. Electrophoresis 23:2592.

25. Fyta, M., S. Melchionna, and S. Succi, 2011. Translocation of biomolecules through solid-state nanopores: Theory meets experiments. Journal of Polymer Science Part B: Polymer Physics 49:985.

26. Muthukumar, M., 2011. Polymer Translocation. CRC Press, Taylor & Francis Group, Boca Raton, FL, USA.

27. Sakaue, T., 2007. Nonequilibrium dynamics of polymer translocation and straightening. Physical Review E 76:021803.

28. Fyta, M., S. Melchionna, S. Succi, and E. Kaxiras, 2008. Hydrodynamic correlations in the translocation of a biopolymer through a nanopore: Theory and multiscale simulations. Physical Review E 78:036704.

29. Lam, P.-M., and Y. Zhen, 2015. Dynamic scaling theory of the forced translocation of a semi-flexible polymer through a nanopore. Journal of Statistical Physics 161:197.

30. Sakaue, T., 2016. Dynamics of polymer translocation: A short review with an introduction of weakly-driven regime. Polymers 8:424.

31. Sarabadani, J., T. Ikonen, H. Mökkönen, T. Ala-Nissila, S. Carson, and M. Wanunu, 2017. Driven translocation of a semi-flexible polymer through a nanopore. Scientific Reports 7:7423.

32. Hsiao, P.-Y., 2018. Translocation of Charged Polymers through a Nanopore in Monovalent and Divalent Salt Solutions: A Scaling Study Exploring over the Entire Driving Force Regimes. Polymers 10:1229.

33. Sung, W., and P. Park, 1996. Polymer translocation through a pore in a membrane. Physical Review Letters 77:783.

34. Ambjörnsson, T., S. P. Apell, Z. Konkoli, E. A. Di Marzio, and J. J. Kasianowicz, 2002. Charged polymer membrane translocation. Journal of Chemical Physics 117:4063.

35. Muthukumar, M., 2003. Polymer escape through a nanopore. Journal of Chemical Physics 118:5174.

36. Kong, C., and M. Muthukumar, 2004. Polymer translocation through a nanopore. II. Excluded volume effect. Journal of Chemical Physics 120:3460.

37. Muthukumar, M., 2007. Mechanism of DNA transport through pores. Annual Review of Biophysics and Biomolecular Structure 36:435.

38. Panja, D., G. T. Barkema, and R. C. Ball, 2007. Anomalous dynamics of unbiased polymer translocation through a narrow pore. Journal of Physics: Condensed Matter 19:432202.

39. Buyukdagli, S., and T. Ala-Nissila, 2016. Electrostatic energy barriers from dielectric membranes upon approach of translocating DNA molecules. Journal of Chemical Physics 144:084902.

40. Zhang, C., X. Lin, and H. Yang, 2016. Theoretical model of biomacromolecule through nanopore including effects of electrolyte and excluded volume. Applied Mathematics and Mechanics 37:787.

41. Buyukdagli, S., J. Sarabadani, and T. Ala-Nissila, 2019. Theoretical modeling of polymer translocation: From the electrohydrodynamics of short polymers to the fluctuating long polymers. Polymers 11:118.

42. Hernández-Ortiz, J. P., M. Chopra, S. Geier, and J. J. de Pablo, 2009. Hydrodynamic effects on the translocation rate of a polymer through a pore. Journal of Chemical Physics 131:044904.

43. Kapahnke, F., U. Schmidt, D. W. Heermann, and M. Weiss, 2010. Polymer translocation through a nanopore: The effect of solvent conditions. Journal of Chemical Physics 132:164904.

44. Moisio, J. E., J. Piili, and R. P. Linna, 2016. Driven polymer translocation in good and bad solvent: Effects of hydrodynamics and tension propagation. Physical Review E 94:022501.

45. Zwanzig, R., 2001. Nonequilibrium Statistical Mechanics. Oxford University Press, USA.

46. Zwanzig, R., 1992. Diffusion past an entropy barrier. Journal of Physical Chemistry 96:3926.

47. M. Doi and S. F. Edwards, 1986. The Theory of Polymer Dynamics. Clarendon, Oxford, UK.

48. Rubinstein, M. C., and R. H. Colby, 2003. Polymer Physics. Oxford University Press, Oxford, UK.

49. Rouse Jr, P. E., 1953. A theory of the linear viscoelastic properties of dilute solutions of coiling polymers. Journal of Chemical Physics 21:1272.

50. Zimm, B. H., 1956. Dynamics of polymer molecules in dilute solution: viscoelasticity, flow birefringence and dielectric loss. Journal of Chemical Physics 24:269.

51. Freed, K. F., 1972. Functional Integrals and Polymer Statistics. Advances in Chemical Physics 22:1.

52. Jackson, J. D., 1998. Classical Electrodynamics. Wiley, NJ, USA.

53. Chandrasekhar, S., 1943. Stochastic problems in physics and astronomy. Reviews of Modern Physics 15:1.

